# Protein structure prediction and design for high-throughput computing

**DOI:** 10.1101/2025.07.18.665594

**Authors:** Vinay Saji Mathew, Gretta D. Kellogg, William KM Lai

## Abstract

Recent advances in structural biology and machines learning have resulted in a revolution in molecular biology. This revolution is driven by protein structure prediction and design tools such as Alphafold3, Chai-1, and Boltz-2 which are now able to accurately model protein structures as well as predict protein-complex formation with a variety of substrates at atomic resolution (i.e., DNA, RNA, small ligands, post-translational modifications). The impact of these protein-structure prediction algorithms has been matched by the emergence of *in silico* protein design platforms (RFdiffusion), which now promise to revolutionize synthetic biology and novel disease therapeutics. Despite their potential to transform molecular biology, the adoption of these algorithms is hindered in part, not only by their high computational requirements, but also by the difficulty in deploying these algorithms on available systems. To help address these barriers, we developed containerized solutions for AlphaFold3, Chai-1, Boltz-2, and RFdiffusion, optimized across a variety of computational architectures (e.g., x86 and ARM). Additionally, we present OmniFold, an optimized wrapper-platform with automatic QC report generation that enables AlphaFold3, Chai-1, and Boltz-2 to perform simultaneously while more efficiently utilizing GPU systems. Precompiled containers and their definition files are available as open source through Sylabs and GitHub. We hope that these containers and repos will help to facilitate reproducibility, accessibility, and accelerate scientific discovery.

**Availability and implementation:** Source code for containers is available at:

https://github.com/EpiGenomicsCode/ProteinStruct-Containers

https://github.com/EpiGenomicsCode/ProteinDesign-Containers

https://github.com/EpiGenomicsCode/OmniFold

## Introduction

The computational capability to accurately predict the 3D structure of a protein has transformed vast swathes of molecular biology [1-5]. This is due in no small part to one of the central tenets in biology: function follows form. The ability to accurately infer form enables researchers to identify functional characteristics of proteins. The problem of predicting 3D protein structure had been long outstanding due to the computational and biochemical challenge inherent in modeling the complex high-dimensional atomic interactions involved in a protein achieving its structure. Simultaneous advances in structural biology (i.e., cyroEM) and applied machine learning and artificial intelligence (i.e., transformers) have produced enough training data and the computational frameworks needed to leverage said data [6, 7]. This fusion of domain-knowledge resulted in a breakthrough (AlphaFold) on the problem of computational protein folding recently described by Google’s DeepMind AI team [3].

Iterations of the AlphaFold algorithm (AlphaFold2-multimer) added the ability to accurately model protein-protein interactions from amino-acid sequence [8]. The most recent version of the algorithm (AlphaFold3) has provided dramatic improvement to its capabilities, including the ability to model proteins with DNA, RNA, small molecules, and others [5]. Similar approaches such as Chai-1 and Boltz-2 have since arisen which offer similar but distinct prediction capabilities relative to AlphaFold3 [9, 10]. One of the more provocative capabilities that have arisen from the rise of AI-driven protein structure prediction has been the ability to design novel proteins completely *in silico* [11, 12]. Pioneered by the Baker lab, RFdiffusion has emerged as the dominant novel protein design platform with biochemical functionality [13, 14]. The ability to design custom proteins with specific biochemical functionality (e.g., immune, enzymatic, drug design) has the potential to overturn entire fields of biochemistry and kickstart the next stage in synthetic biology.

The adoption and availability of these algorithms has been rate-limited by the fact that most of these protein-structure prediction algorithms are particularly resource-intensive, requiring substantial CPU, GPU, memory, storage, and I/O bandwidth to perform at scale. Broadly speaking, these algorithms typically rely on a genetic search using multiple sequence alignment algorithms (MSAs) against extensive databases of genomic sequence. These MSAs are then used as a weighted input to a trained neural network that assembles and accurately models the 3D structure of proteins. The user is then presented with a predicted structure at atomic resolution and resulting quality control metrics.

Given the shared-access nature of many public HPC systems, these computational requirements can pose significant challenges to institutions looking to offer these algorithms as a supported service for tens to hundreds of concurrent researchers [15]. To promote broader adoption of these algorithms, we have developed containerized solutions for AlphaFold3, Chai-1, Boltz-2, and RFDiffusion designed to operate on a variety of underlying CPU and GPU architectures. Crucially, in support of optimal GPU resource utilization, we also developed OmniFold, a protein ensemble approach, that enables simultaneous running of AlphaFold3, Chai-1, and Bolt-2 from the same source MSAs. OmniFold performs automatic quality control analysis after inference and assembles the results into in a simple HTML file that provides a central point of interpretation for all the structures generated. We provide precompiled containers ready to use and freely available on Sylabs as well as the singularity definition files with build and usage instructions on GitHub. It is our hope that these resources will provide increased accessibility for these groundbreaking algorithms to the broader community.

## Data availability

### Sylabs containers

1. AlphaFold 3 (x86/ARM): https://cloud.sylabs.io/library/vinaymatt/repo/alphafold3
2. Boltz - ARM: https://cloud.sylabs.io/library/boltzarm/repo/boltzarm
3. Boltz – x86: https://cloud.sylabs.io/library/boltzx86/repo/boltzx86
4. Chai-1 ARM: https://cloud.sylabs.io/library/chai_labarm
5. Chai-1 x86: https://cloud.sylabs.io/library/chaix86

### Singularity definition files and implementation

- https://github.com/EpiGenomicsCode/ProteinStruct-Containers
- https://github.com/EpiGenomicsCode/ProteinDesign-Containers
- https://github.com/EpiGenomicsCode/OmniFold

## Implementation

### Overview

We provide Singularity containers for three state-of-the-art structure-prediction models: AlphaFold 3 (AF3), Boltz-2, and Chai-1 and one protein-design model: RFdiffusion (**Figure 1**). All build recipes are version-locked and tested on Ubuntu 22.04 systems with NVIDIA GPUs. Pre-built images for all models are available via Sylabs. We also supply definition files and minimal launcher scripts so that the containers can be rebuilt reproducibly on both x86-64 and ARM64 hardware in support of FAIR compliance [16].

**Figure 1.**
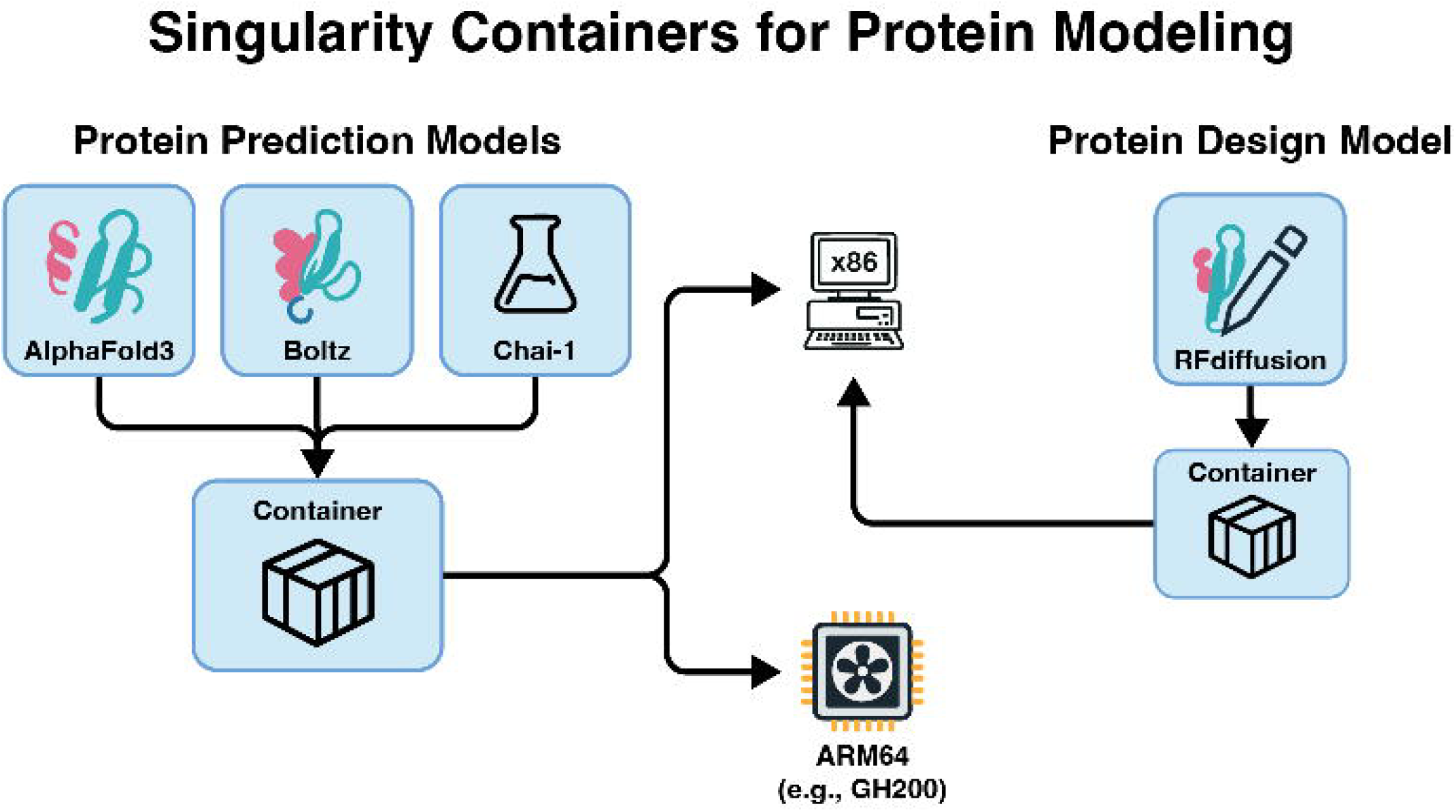
Containers for protein structure prediction and design.

### Pre-built containers

We provide all containers on Sylabs for immediate download. We recommend verifying the authenticity of the container:

~~~
singularity key import keys/mypublic.pem
~~~

Individual containers can be downloaded directly to any system with singularity/apptainer installed with the following commands:

AlphaFold 3 (ARM64 architecture)

~~~
singularity pull --arch arm64 library://vinaymatt/repo/alphafold3:arm64
~~~

AlphaFold 3 (x86_64 architecture)

~~~
singularity pull --arch amd64 library://vinaymatt/repo/alphafold3:amd64
~~~

Boltz-2 (ARM64 architecture)

~~~
singularity pull --arch arm64 library://boltzarm/repo/boltzarm:arm64
~~~

Boltz-2 (x86_64 architecture)

~~~
singularity pull --arch amd64 library://boltzx86/repo/boltzx86:latest
~~~

Chai-1 (ARM64 architecture)

~~~
singularity pull --arch arm64 library://chai_labarm/repo/chai_lab:arm64
~~~

Chai-1 (x86_64 architecture)

~~~
singularity pull --arch amd64 library://chaix86/repo/chai_lab:amd64
~~~

### Cross-architecture build strategy

The x86 builds leverage existing pre-compiled Python wheels served through PyPI or the NVIDIA PyTorch channel. Containers operating on ARM architecture require additional configuration as many core dependencies have no official aarch64 wheels (e.g., Triton, JAX, PyTorch, and certain CUDA tooling). Therefore, our ARM definition files require building these libraries from source with CMake or the package’s own build system. To maximize reproducibility, we have pinned commit hashes to known validated versions (**Table 1**).

**Table 1.** Summary of architecture-specific build decisions.

| Model | Architecture | Source-built components | Pre-staged assets |
| --- | --- | --- | --- |
| AF3 | x86-64 | — (binary wheels) | None |
| AF3 | ARM64 | Triton 3.1, JAX 0.4.34, HMMER | — |
| Boltz-1 | x86-64 | — | ccd.pkl, boltz1_conf.ckpt |
| Boltz-1 | ARM64 | PyTorch 2.7, Triton 3.3 via pip test index | Same as x86 |
| Chai-1 | x86-64 | — | model_v2 weights, ESM embeddings |
| Chai-1 | ARM64 | PyTorch 2.4 built from source for SM 8.9 | Same as x86 |
| RFdiffusion | x86-64 only | — | weights downloaded in post |

The command to compile each protein prediction container is as follows:

~~~
sbatch build_container.slurm <model_directory_name> <architecture>
/path/to/your/build/directory
~~~

Where target architecture can be either ‘`x86’` or ‘`arm’`.

### Runtime wrappers

Each container registers a minimal runscript that activates the environment and executes the model’s Python entrypoint. For standard usage, we recommend using the provided launcher scripts, which abstract over the singularity exec syntax, enforce --nv, and apply sensible defaults for GPU visibility, pre-allocation, and output paths.

Each singularity container can be activated using the common generalized command shown:

~~~
singularity exec --nv/path/to/your/<model >_<arch>.sif
<command_specific_to_model> <arguments>
~~~

Where `<command_specific_to_model>` follows the example launcher commands detailed below.

### AlphaFold 3

AF3 is installed directly from the official repository for x86, augmenting the runtime with CUDA 12.6 wheels for JAX-Triton. The ARM image shares the same base CUDA stack but compiles Triton and JAX from source, as binary wheels are not available. HMMER is built to support the MSA pipeline and mounted into $PATH. A lightweight launcher (run_alphafold3_launcher.py) wraps singularity exec, binds the databases, model weights, and common JAX/XLA environment variables.

Example of general execution:

~~~
python alphafold3/run_alphafold3_launcher.py --json_path=/path/to/input.json -
-model_dir=/path/to/model_params --db_dir=/path/to/databases --
output_dir=/path/to/output [--other-flags…]
~~~

Detailed flags for the Alphafold3 container are as follows:

~~~
--json_path: Path to a single input JSON file.
--input_dir: Path to a directory of input JSON files (alternative to --json_path).
--model_dir: Path to the downloaded AlphaFold 3 model parameters.
--db_dir: Path(s) to the downloaded databases (can be specified multiple times).
--output_dir: Directory where results will be saved.
--use_gpu: Set to false to run without GPU (only data pipeline).
--run_data_pipeline=false: Skip the data pipeline step.
--run_inference=false: Skip the inference step.
~~~

Users may also run `python alphafold3/run_alphafold3_launcher.py --help` to see all available options.

### Boltz-2

Boltz ships binary wheels for x86, so the container installs the package as well as PyTorch 2.7 linked against CUDA 12.8. For ARM compatibility, we bootstrap Miniforge, create an isolated conda environment, and pull test-index wheels for the same PyTorch/Triton versions. Model checkpoints (ccd.pkl and boltz1_conf.ckpt) are cached under a cache folder and exposed through the BOLTZ_CACHE variable. The launcher script we provide resolves the correct image and handles optional ColabFold MSA generation.

Example of general execution:

~~~
python boltz/run_boltz_launcher.py --input_data=/path/to/your/input.fasta --
out_dir=/path/to/output [--other-flags…]
~~~

Detailed flags for the Boltz-2 container are as follows:

~~~
--sif_path: Path to the Boltz Singularity image SIF file (overrides environment
variable and hardcoded path).
--input_data: (Required) Input data file or directory (FASTA/YAML).
--out_dir: Output directory for predictions.
--boltz_cache_dir: Directory for Boltz to download data/models (defaults to
$BOLTZ_CACHE or ∼/.boltz).
--checkpoint: Optional path to a model checkpoint file.
--use_gpu: Enable NVIDIA runtime for GPU usage (default: True).
--gpu_devices: Comma-separated list of GPU devices for
NVIDIA_VISIBLE_DEVICES.
~~~

Users may run `python boltz/run_boltz_launcher.py --help` to see all available options. Additional options include additional Boltz parameters such as `--recycling_steps, -- sampling_steps, --use_msa_server,` etc.

### Chai-1

Chai-1 relies on PyTorch 2.4 kernels that are not yet published for ARM. As a result, the ARM definition stage builds PyTorch from source with TORCH_CUDA_ARCH_LIST=9.0 to target functionality on GH200 Grace Hopper GPUs. Exact versioning should be modified based on the environment the user is working with. Weights and ESM embeddings are downloaded into a downloads folder in both images. As with the other algorithms, the launcher exposes the directory variable and forwards all remaining CLI arguments verbatim to chai-lab.

Example of general execution:

~~~
python chai_1/run_chailab_launcher.py --
fasta_file=/path/to/your/sequence.fasta --output_dir=/path/to/chai_output [--
other-flags…]
~~~

Detailed flags for the Chai-1 container are as follows:

~~~
--sif_path: Path to the Chai Lab Singularity image SIF file (overrides environment
variable and hardcoded path).
--fasta_file: (Required) Path to the input FASTA file.
--output_dir: Path to a directory for storing results.
--force_output_dir: If True, use the exact output directory even if non-empty.
--use_gpu: Enable NVIDIA runtime for GPU usage (default: True).
--gpu_devices: Comma-separated list of GPU devices for
NVIDIA_VISIBLE_DEVICES.
~~~

Users may run `python chai_1/run_chailab_launcher.py --help` to see all available options. Additional options for Chai-1 features include `--msa_directory, --constraint_path, -- num_diffn_samples`, etc.

### OmniFold

We developed OmniFold to provide a unified protein prediction ensemble framework for running multiple sequence prediction algorithms in parallel. OmniFold currently supports the simultaneous operation of AlphaFold3, Chai-1, and Boltz-2. OmniFold will run each model (e.g., AlphaFold3, Boltz, Chai-1) based on whether their respective Singularity image paths (−- alphafold3_sif_path, --boltz1_sif_path, --chai1_sif_path) have been provided by the user. Models without a SIF path are skipped.

We architected OmniFold to be extensible to simultaneously process any number of prediction algorithms while minimizing any additional computational overhead inherent to running multiple MSA generation scripts. Specifically, we have developed a unified input handling process that parses a single input file (e.g., FASTA, AF3 JSON, or Boltz YAML) and standardizes the sequence and chain information internally.

OmniFold first determines if MSAs are needed based on your input and --msa_method flag. If the selected MSA method is AlphaFold3 (default), OmniFold will initially run the AlphaFold3 data pipeline through the Singularity container. We developed modified versions of AlphaFold3’s internal pipeline.py and run_alphafold.py scripts to ensure compatible A3M file generation (e.g., UniRef90, MGnify, etc.) for each chain within the standard AlphaFold3 output structure. MSA caching is restructured to enable this change. The resulting AlphaFold3 MSA JSON is then parsed to extract per-protein A3M files that are then made available for Boltz. If Chai-1 is also set to be run, the A3M files from the AlphaFold3 output (msas/chain_X/*.a3m) are converted into Chai-1’s PQT format (msas_forChai/*.pqt).

OmniFold also directly supports ColabFold-generated MSAs [17]. If the user-specified MSA method is ‘ColabFold’, OmniFold will directly query the ColabFold API to retrieve MSAs. The resulting A3M files are piped directly to the AlphaFold3 pipeline, while Boltz and Chai-1 consume the same cached MSAs using their native integrations. As with the stock versions of the algorithms, OmniFold also supports using existing MSAs from the input files to bypass their calculation.

OmniFold detects available GPUs and will assign each model to an available GPU. When run on a Grace Hopper GH200 superchip, OmniFold will run all 3 models in parallel on the same chip. Alternatively, if only a single GPU is made available, OmniFold will run each model sequentially, reusing the same MSAs for each model.

The output produced by all 3 inference models working in parallel can be substantial, especially for large multimeric complexes, relative to running the models sequentially. OmniFold organizes all pertinent results and logs from all executed model runs into the specified output directory. Additionally, we developed a custom report generation script that automatically generates a quality control report comparing the resulting structures across algorithms (**Figure 2A**). This report is consolidated into a single HTML file making it viewable on all modern web browsers. We calculate and display key metrics (i.e., pLDDT, ipSAE, pDockQ) (**Figure 2B,C**) [18-20]. We also modified the PAE viewer platform to natively visualize Boltz-2 and Chai-1 derived structures and bundled it into our OmniFold report platform [21].

**Figure 2.**
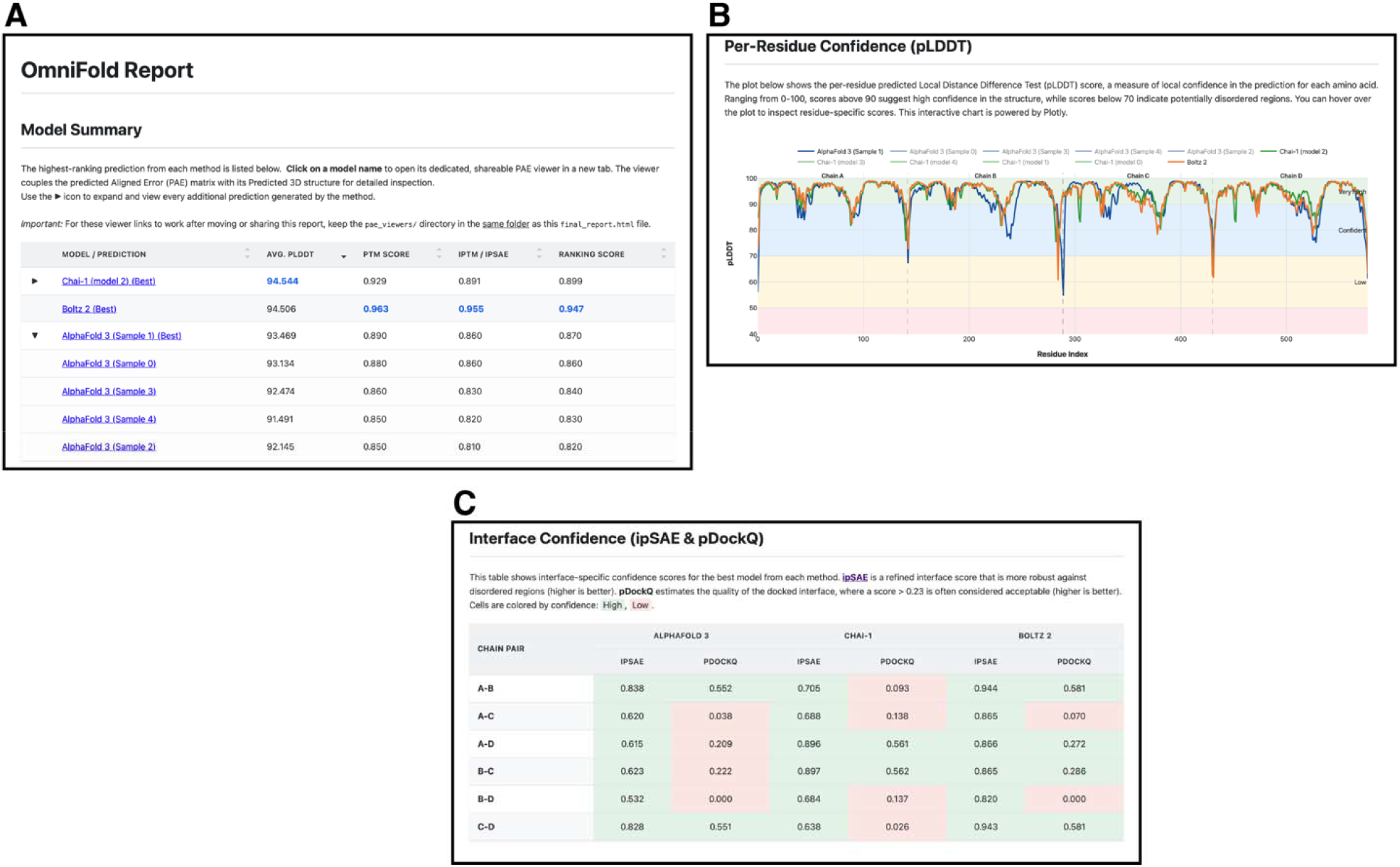
**A**. OmniFold quality control report with sortable scores for each structure generated by all available protein prediction engines. B. Interactive plot displaying the per-residue pLDDT score for each generated structure. C. Interface confidence scores (ipSAE and pDockQ) for multimeric predictions.

### RFdiffusion

Our container pins NVIDIA CUDA 11.6.2 and pulls torch-1.12 and DGL-1.0 wheels compiled for that toolchain. The official download script runs during post and installs the weights into rfdiffusion.models so that no external volume is required at runtime. A single-file launcher mirrors the pattern used for the prediction models.

Command to compile the container:

~~~
sbatch build_container.slurm rfdiffusion x86 /path/to/your/build/directory
~~~

Example of general execution:

~~~
singularity exec --nv /path/to/your/rfdiffusion_x86.sif
inference.output_prefix=/outputs/test_design
inference.input_pdb=/inputs/my_protein.pdb ‘contigmap.contigs=[100-100]’
inference.num_designs=1
~~~

Detailed flags for the RFdiffusion container are as follows:

~~~
--sif_path: Path to the RFdiffusion Singularity image SIF file (overrides environment
variable and hardcoded path).
--input_pdb: Path to the input PDB file (e.g., for motif scaffolding, binder design).
Corresponds to inference.input_pdb.
--output_prefix: (Required) Prefix for output files. Corresponds to
inference.output_prefix.
--model_directory_path: Path to custom model weights if not using the
container’s default. Corresponds to inference.model_directory_path.
--num_designs: Number of designs to generate. Corresponds to
inference.num_designs.
--use_gpu: Set to true (default) or false to enable/disable NVIDIA GPU runtime.
--gpu_devices: Comma-separated list of GPU devices for
NVIDIA_VISIBLE_DEVICES (e.g., ‘0,1’).
--bind_mounts: List of additional paths to bind mount (e.g.,
“/host/data:/data_in_container”).
~~~

RFdiffusion specific commands are extensive and the primary RFdiffusion GitHub repository from should be consulted for further details (https://github.com/RosettaCommons/RFdiffusion).

### Databases and model weights

Databases for MSA can be downloaded following each algorithm’s specific recommendation. Model weights for Alphafold3 are restricted by Google and require explicit registration to acquire and use. For large-scale deployments, DeepMind provides a non-commercial research institution Terms of Use option for deploying centralized weights through an institutional-wide license. Model weights for Boltz-2, Chai-1, and RFdiffusion are included directly within the container and do not need to be downloaded separately.

### Systems hardware

All software developed here used NCSA’s Delta and DeltaAI systems provided through NSF-ACCESS. Exact system specifications:

- x86 Arch with H200
  - 2 × Intel Xeon Platinum 8558 (48 cores @ 2.1 GHz) → 96 CPU cores total
  - 1 × NVIDIA H200 SXM (144 GB HBM3e)
  - PCIe 4.0 ×16 CPU ↔ GPU link (∼32 GB s □ ^1^)
  - 1 TB DDR5-5600 system RAM
- ARM Arch on GH200 Superchip
  - 4 × NVIDIA Grace Hopper GH200 modules, each containing
  - 72-core Grace CPU (3.5 GHz, SVE2-512)
  - Hopper-100 GPU (120 GB HBM3e)
  - NVLink-C2C CPU ↔ GPU fabric (∼900 GB s □ ^1^ per module)
  - 288 CPU cores total across the node
  - 480 GB LPDDR5X CPU memory + 480 GB HBM3e GPU memory

## Discussion

We expect that the continued rapid adoption of these protein structure prediction and design algorithms will continue across the broad domain of life sciences. Already, a sizable number of nationally recognized HPC’s (e.g., NIH BioWulf, PSU ROAR, PSC, TACC, etc.) now offer Alphafold3 and companion algorithms to their user-base. The extreme resource intensiveness of these algorithms has made it a challenge for these centers to offer them to the broader community at scale as well as for individual researchers to explore these algorithms in smaller settings [15]. To assist the community, we first developed containerized solutions for AlphaFold3, Chai-1, Boltz-2, and RFDiffusion. These containers were architected to operate on a variety of CPU and GPU architectures to maximize their interoperability across diverse environments.

We recognized that the MSA generation component was often a substantial drain on time and furthermore that the MSAs generated were often cross-compatible across algorithms with minimal parsing. We therefore developed the OmniFold ensemble approach which efficiently recycles MSAs across algorithms and performs load balancing across all available GPUs within the environment. The centralized HTML report generated by OmniFold is an interactive system that compiles and present useful quality metrics for users to further interpret their generated structures.

We envision that as further protein prediction algorithms become available, they will be added into the OmniFold framework where possible. Furthermore, we see an opportunity for better integration between OmniFold and RFdiffusion to make a fully end-to-end protein design and prediction framework that makes optimal use of all available resources.

All our precompiled containers are open source and accessible on Sylabs. To combat ever-present software rot, we have also made all singularity definition files with build and usage instructions available on GitHub. It is our hope that these resources will provide increased accessibility for these groundbreaking algorithms and encourage broader participation by the research community. This work improves upon the architecture and resource commitment required, assisting the national HPC community by enabling easier adoption of this technology in a scalable, rigorous, and reproducible manner.

## Acknowledgements

We thank the members of the Kumara and Lai labs for their comments and feedback. We also thank Matt Hansen and Chad Bahrmann (PSU ICDS) and Charles Pavloski (NCSA NSF-OAC2320345 & NSF-OAC2005572), for their feedback and troubleshooting.

## Funding

This project was supported by NIH-R35GM155380 to W.K.M.L., the Cornell University BRC Epigenomics Core Facility (RRID:SCR_021287), Penn State Institute for Computational and Data Sciences (RRID:SCR_025154), Penn State University Center for Applications of Artificial Intelligence and Machine Learning to Industry Core Facility (RRID:SCR_022867), and a gift to AIMI research from Dell Technologies. Computational support was provided by NSF ACCESS (BIO230041) to W.K.M.L. and G.D.K.

## Notes

https://github.com/EpiGenomicsCode/OmniFold

https://github.com/EpiGenomicsCode/ProteinStruct-Containers

https://github.com/EpiGenomicsCode/ProteinDesign-Containers

### Competing Interest Statement

The authors have declared no competing interest.

## Bibliography

1. Perrakis A, Sixma TK. AI revolutions in biology: The joys and perils of AlphaFold. EMBO Rep. 2021;22(11):e54046. Epub 2021/10/21. doi: 10.15252/embr.202154046. PubMed PMID: 34668287; PubMed Central PMCID: PMC8567224.

2. Thornton JM, Laskowski RA, Borkakoti N. AlphaFold heralds a data-driven revolution in biology and medicine. Nat Med. 2021;27(10):1666–9. Epub 2021/10/14. doi: 10.1038/s41591-021-01533-0. PubMed PMID: 34642488.

3. Jumper J, Evans R, Pritzel A, Green T, Figurnov M, Ronneberger O, et al. Highly accurate protein structure prediction with AlphaFold. Nature. 2021;596(7873):583–9. Epub 2021/07/16. doi: 10.1038/s41586-021-03819-2. PubMed PMID: 34265844.

4. Baek M, DiMaio F, Anishchenko I, Dauparas J, Ovchinnikov S, Lee GR, et al. Accurate prediction of protein structures and interactions using a three-track neural network. Science. 2021;373(6557):871–6. Epub 2021/07/21. doi: 10.1126/science.abj8754. PubMed PMID: 34282049; PubMed Central PMCID: PMC7612213.

5. Abramson J, Adler J, Dunger J, Evans R, Green T, Pritzel A, et al. Accurate structure prediction of biomolecular interactions with AlphaFold 3. Nature. 2024;630(8016):493–500. Epub 2024/05/09. doi: 10.1038/s41586-024-07487-w. PubMed PMID: 38718835.

6. Vaswani A, Shazeer N, Parmar N, Uszkoreit J, Jones L, Gomez AN, et al. Attention Is All You Need2017 June 01, 2017:[1706.03762 p.]. Available from: https://ui.adsabs.harvard.edu/abs/2017arXiv170603762V.

7. Renaud JP, Chari A, Ciferri C, Liu WT, Remigy HW, Stark H, et al. Cryo-EM in drug discovery: achievements, limitations and prospects. Nat Rev Drug Discov. 2018;17(7):471–92. Epub 2018/06/09. doi: 10.1038/nrd.2018.77. PubMed PMID: 29880918.

8. Evans R, O’Neill M, Pritzel A, Antropova N, Senior A, Green T, et al. Protein complex prediction with AlphaFold-Multimer. bioRxiv. 2022:2021.10.04.463034. doi: 10.1101/2021.10.04.463034.

9. team CD, Boitreaud J, Dent J, McPartlon M, Meier J, Reis V, et al. Chai-1: Decoding the molecular interactions of life. bioRxiv. 2024:2024.10.10.615955. doi: 10.1101/2024.10.10.615955.

10. Passaro S, Corso G, Wohlwend J, Reveiz M, Thaler S, Somnath VR, et al. Boltz-2: Towards Accurate and Efficient Binding Affinity Prediction. bioRxiv. 2025:2025.06.14.659707. doi: 10.1101/2025.06.14.659707.

11. Dauparas J, Anishchenko I, Bennett N, Bai H, Ragotte RJ, Milles LF, et al. Robust deep learning-based protein sequence design using ProteinMPNN. Science. 2022;378(6615):49–56. Epub 2022/09/16. doi: 10.1126/science.add2187. PubMed PMID: 36108050; PubMed Central PMCID: PMC9997061.

12. Watson JL, Juergens D, Bennett NR, Trippe BL, Yim J, Eisenach HE, et al. De novo design of protein structure and function with RFdiffusion. Nature. 2023;620(7976):1089–100. Epub 2023/07/12. doi: 10.1038/s41586-023-06415-8. PubMed PMID: 37433327; PubMed Central PMCID: PMC10468394.

13. Vazquez Torres S, Leung PJY, Venkatesh P, Lutz ID, Hink F, Huynh HH, et al. De novo design of high-affinity binders of bioactive helical peptides. Nature. 2024;626(7998):435–42. Epub 2023/12/19. doi: 10.1038/s41586-023-06953-1. PubMed PMID: 38109936; PubMed Central PMCID: PMC10849960 J.L.W., M.J.M., N.R.B. and G.R.L. are inventors on a provisional patent application submitted by the University of Washington for the design and composition of the proteins created in this study.

14. Liu C, Wu K, Choi H, Han H, Zhang X, Watson JL, et al. Diffusing protein binders to intrinsically disordered proteins. bioRxiv. 2024. Epub 2024/07/29. doi: 10.1101/2024.07.16.603789. PubMed PMID: 39071267; PubMed Central PMCID: PMC11275890.

15. Park H, Patel P, Haas R, Huerta EA. APACE: AlphaFold2 and advanced computing as a service for accelerated discovery in biophysics. Proc Natl Acad Sci U S A. 2024;121(27):e2311888121. Epub 20240624. doi: 10.1073/pnas.2311888121. PubMed PMID: 38913887; PubMed Central PMCID: PMC11228474.

16. Wilkinson MD, Dumontier M, Aalbersberg IJ, Appleton G, Axton M, Baak A, et al. The FAIR Guiding Principles for scientific data management and stewardship. Sci Data. 2016;3:160018. Epub 2016/03/16. doi: 10.1038/sdata.2016.18. PubMed PMID: 26978244; PubMed Central PMCID: PMC4792175 Honorary Academic Editor and consultant.

17. Mirdita M, Schutze K, Moriwaki Y, Heo L, Ovchinnikov S, Steinegger M. ColabFold: making protein folding accessible to all. Nat Methods. 2022;19(6):679–82. Epub 20220530. doi: 10.1038/s41592-022-01488-1. PubMed PMID: 35637307; PubMed Central PMCID: PMC9184281.

18. Basu S, Wallner B. DockQ: A Quality Measure for Protein-Protein Docking Models. PLoS One. 2016;11(8):e0161879. Epub 2016/08/26. doi: 10.1371/journal.pone.0161879. PubMed PMID: 27560519; PubMed Central PMCID: PMC4999177.

19. Bryant P, Pozzati G, Elofsson A. Improved prediction of protein-protein interactions using AlphaFold2. Nat Commun. 2022;13(1):1265. Epub 20220310. doi: 10.1038/s41467-022-28865-w. PubMed PMID: 35273146; PubMed Central PMCID: PMC8913741.

20. Dunbrack RL. Rēs ipSAE loquunt: What’s wrong with AlphaFold’s ipTM score and how to fix it. bioRxiv. 2025:2025.02.10.637595. doi: 10.1101/2025.02.10.637595.

21. Elfmann C, Stulke J. PAE viewer: a webserver for the interactive visualization of the predicted aligned error for multimer structure predictions and crosslinks. Nucleic Acids Res. 2023;51(W1):W404–W10. doi: 10.1093/nar/gkad350. PubMed PMID: 37140053; PubMed Central PMCID: PMC10320053.

